## Supplemental Materials for "Biomolecular condensates form spatially inhomogeneous network fluids"

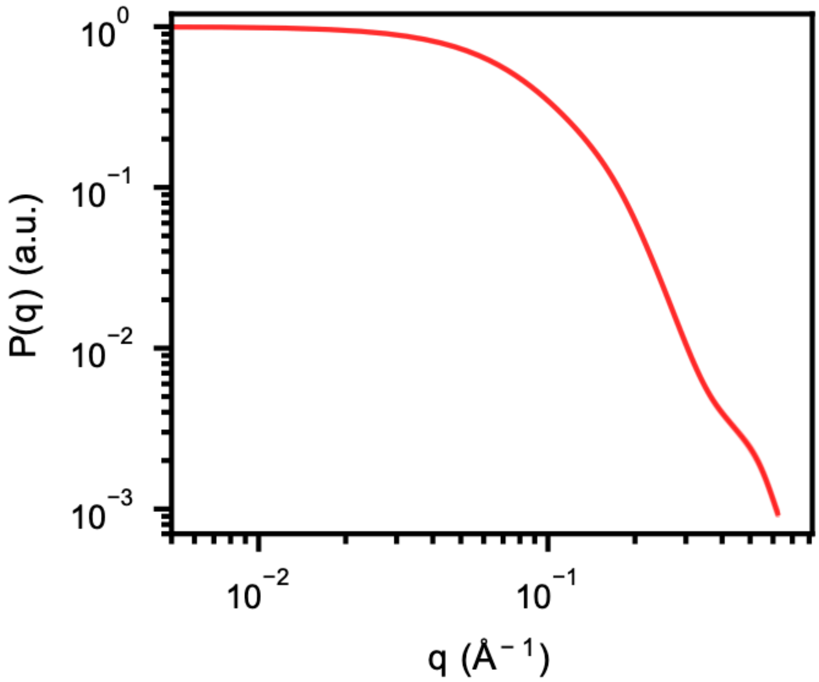


**Supplementary Fig. S1: Scattering probability using CAMPARI for N130 pentamers.** Snapshots from ABSINTH-based atomistic simulations of N130 in the presence of rpL5 peptides were culled, and the scattering profiles were computed for N130 pentamers, whilst excluding the rpL5 peptides. The N130 molecules include the oligomerization domains and disordered regions (see Fig. 1a in the main text). Details of how the scattering profiles were computed are as described in the methods section.

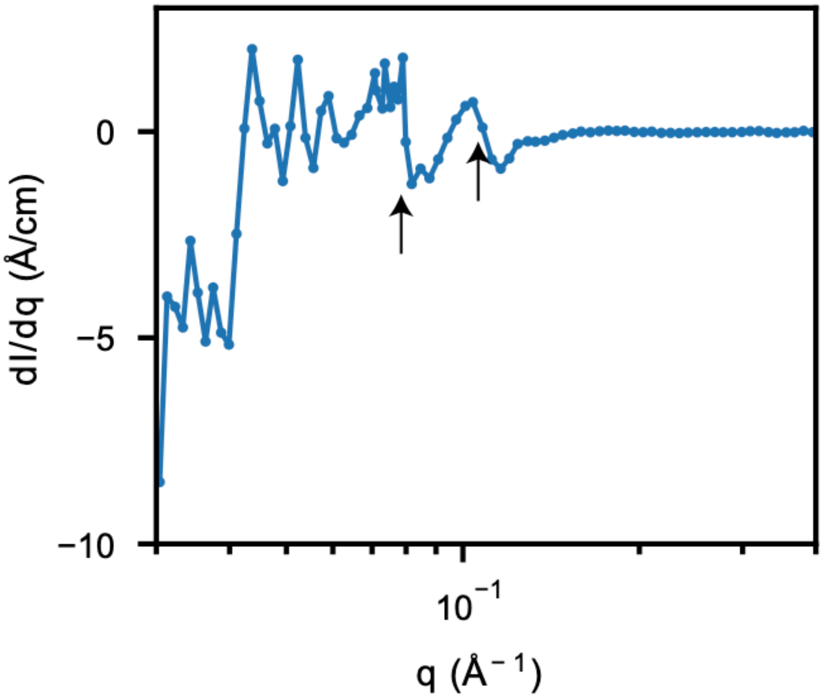


**Supplementary Fig. S2: Derivative analysis of the SANS curve for N130+rpL5.** The first derivative, $dI/dq$, of the SANS data confirms the assignment of the two peak positions at large-$q$ corresponding to ~55 Å (right arrow) and ~77 Å (left arrow). Due to low signal-to-noise, unambiguous assignment of additional peaks is not possible using this analysis.

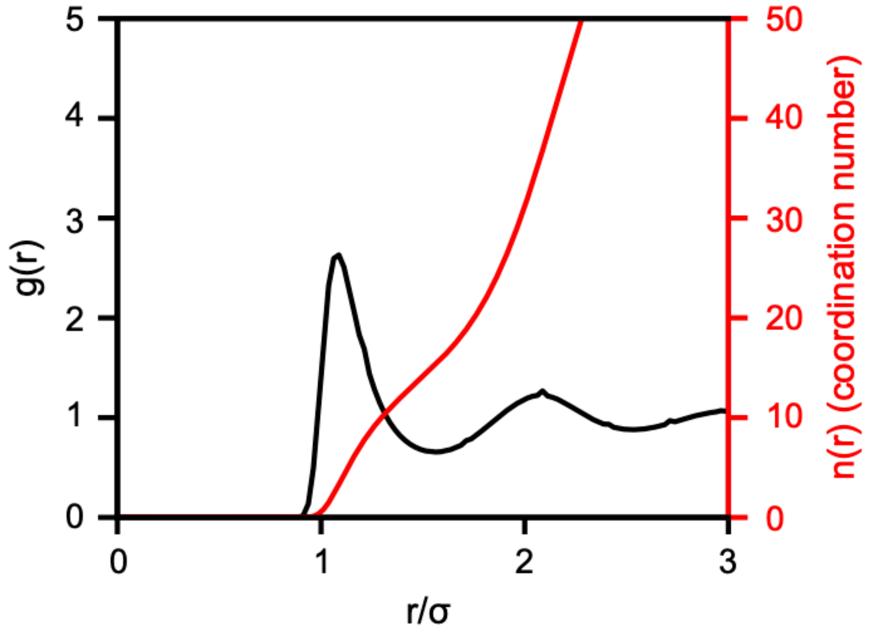


**Supplementary Fig. S3: Radial distribution function of a simple liquid.** Radial distribution function $g(r)$ showing the correlations between particles in a Lennard-Jones fluid along with the volume integral $n(r)$ quantifying the mean coordination number as a function of interparticle separation $r$. On average, approximately twelve Lennard-Jones particles are found within the first coordination shell.

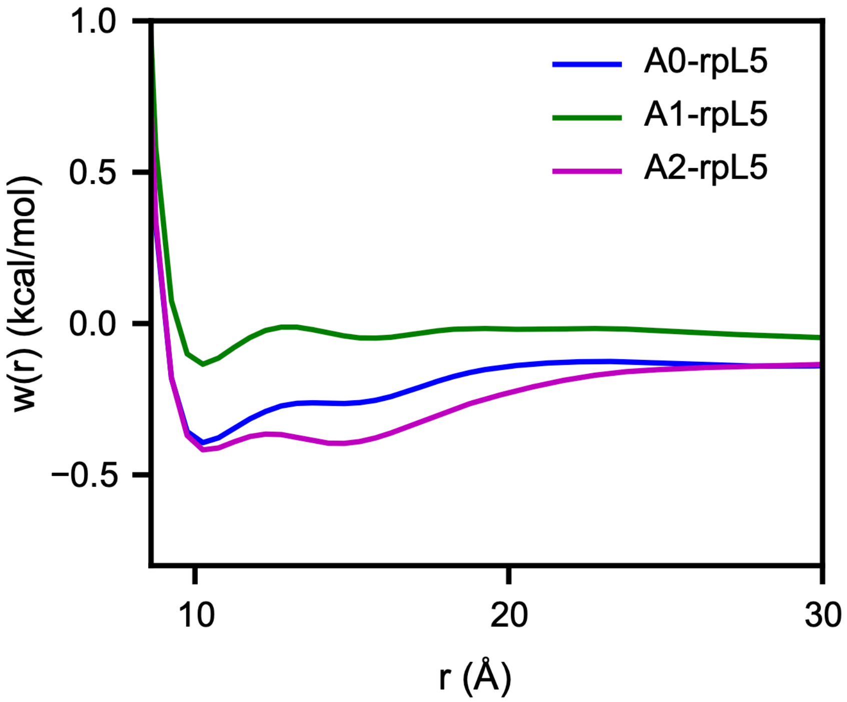


**Supplementary Fig. S4: Potentials of mean force calculated with respect to the correlations between the negative charges in the acidic tracts of N130 and the positive charges in rpL5.** The potential of mean force was calculated using $w(r) = -RT\ln g(r)$, where $R$ is the gas constant, and $T$ is the simulation temperature. The trend whereby the depths of the first minima follow the pattern A2 < A0 < A1 indicates the interaction hierarchy among the tracts as evident in the corresponding $g(r)$ and the contact map.

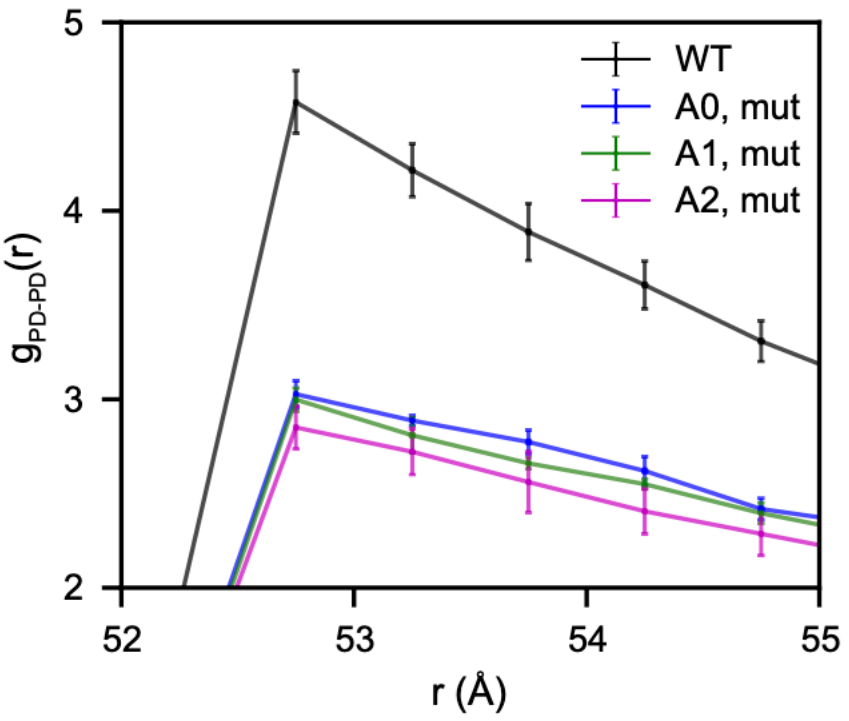


**Supplementary Fig. S5: The reduction in the first maximum of $g_{PD\text{-}PD}(r)$ computed after neutralizing the charges in select acidic tracts.** The magnitude of the reduction is greatest for the neutralized A2, followed by the A0 and A1 mutants. The values of the peak heights of the A0 and A1 mutants are statistically similar within error. The error bars represent the standard error about the mean over five replicates.

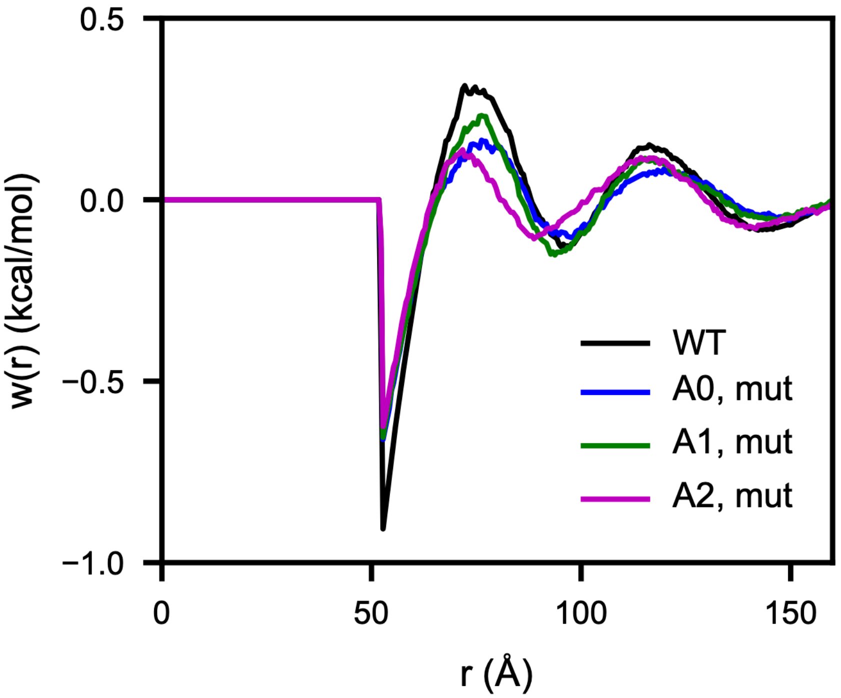


**Supplementary Fig. S6: Potentials of mean force calculated with respect to the PD after neutralizing the charges in select acidic tracts.** The potential of mean force was calculated as before according to $w(r) = -RT\ln g(r)$, where $R$ is the gas constant, and $T$ is the simulation temperature. The trend at $r{\sim}53$ Å whereby $w(r)$ is less favorable for all acidic tracts compared to wild type, and A2 has the smallest value is also reflected in the corresponding $g(r)$ and the contact map.

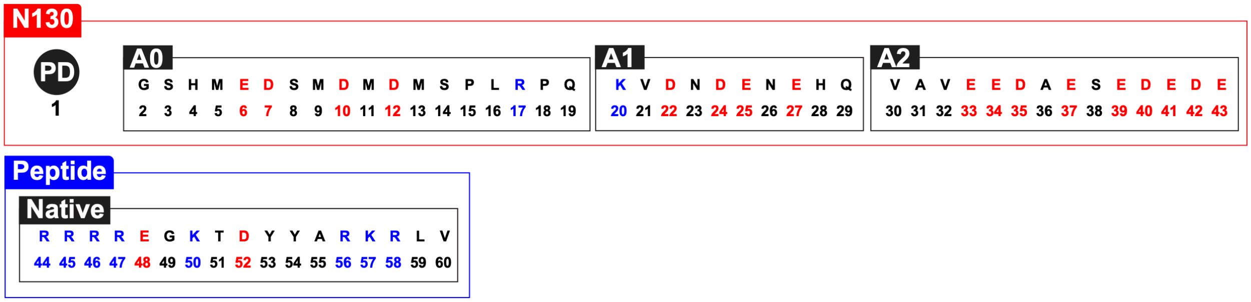


**Supplementary Fig. S7: Bead types in the coarse-grained model.** The bead types and corresponding amino acid residues are given for the pentamerized domain (PD) and acidic tracts of N130 and the native peptide.

**Supplementary Table S1:** LJ parameters for the native rpL5 peptide.

| Type 1 | Type 2 | $\epsilon$ (kcal/mol) | $\sigma$ (Å) | $r_c$ (Å) |
|---|---|---|---|---|
| 1 | * | 100.0 | 26.16 | 26.16 |
| 1 | 1 | 100.0 | 52.32 | 52.32 |
| 2 | 2 | 0.25 | 18.66 | 46.66 |
| 3 | 3 | 0.25 | 3.65 | 9.11 |
| 4 | 4 | 0.73 | 5.02 | 12.54 |
| 5 | 5 | 0.73 | 5.95 | 14.88 |
| 6 | 6 | 0.2 | 10.37 | 25.92 |
| 7 | 7 | 0.2 | 10.37 | 25.92 |
| 8 | 8 | 0.25 | 3.61 | 9.01 |
| 9 | 9 | 0.73 | 5.07 | 12.67 |
| 10 | 10 | 0.2 | 10.37 | 25.92 |
| 11 | 11 | 0.73 | 5.10 | 12.74 |
| 12 | 12 | 0.2 | 10.37 | 25.92 |
| 13 | 13 | 0.73 | 5.05 | 12.63 |
| 14 | 14 | 0.25 | 3.60 | 8.99 |
| 15 | 15 | 0.25 | 3.86 | 9.64 |
| 16 | 16 | 0.73 | 4.60 | 11.49 |
| 17 | 17 | 0.2 | 10.37 | 25.92 |
| 18 | 18 | 0.25 | 3.85 | 9.62 |
| 19 | 19 | 0.73 | 4.69 | 11.72 |
| 20 | 20 | 0.2 | 10.37 | 25.92 |
| 21 | 21 | 0.73 | 4.08 | 10.21 |
| 22 | 22 | 0.2 | 10.37 | 25.92 |
| 23 | 23 | 0.73 | 4.20 | 10.50 |
| 24 | 24 | 0.2 | 10.37 | 25.92 |
| 25 | 25 | 0.2 | 10.37 | 25.92 |
| 26 | 26 | 0.73 | 4.24 | 10.59 |
| 27 | 27 | 0.2 | 10.37 | 25.92 |
| 28 | 28 | 0.73 | 4.99 | 12.46 |
| 29 | 29 | 0.73 | 4.67 | 11.69 |
| 30 | 30 | 0.73 | 4.08 | 10.19 |
| 31 | 31 | 0.25 | 3.39 | 8.47 |
| 32 | 32 | 0.73 | 4.08 | 10.20 |
| 33 | 33 | 0.2 | 10.37 | 25.92 |
| 34 | 34 | 0.2 | 10.37 | 25.92 |
| 35 | 35 | 0.2 | 10.37 | 25.92 |
| 36 | 36 | 0.25 | 3.39 | 8.47 |
| 37 | 37 | 0.2 | 10.37 | 25.92 |
| 38 | 38 | 0.25 | 3.61 | 9.04 |
| 39 | 39 | 0.2 | 10.37 | 25.92 |
| 40 | 40 | 0.2 | 10.37 | 25.92 |
| 41 | 41 | 0.2 | 10.37 | 25.92 |

| | | | | |
|---|---|---|---|---|
| 42 | 42 | 0.2 | 10.37 | 25.92 |
| 43 | 43 | 0.2 | 10.37 | 25.92 |
| 44 | 44 | 0.2 | 8.5 | 21.25 |
| 45 | 45 | 0.2 | 8.5 | 21.25 |
| 46 | 46 | 0.2 | 8.5 | 21.25 |
| 47 | 47 | 0.2 | 8.5 | 21.25 |
| 48 | 48 | 0.2 | 8.5 | 21.25 |
| 49 | 49 | 0.01 | 2.96 | 7.39 |
| 50 | 50 | 0.2 | 8.5 | 21.25 |
| 51 | 51 | 0.01 | 3.97 | 9.92 |
| 52 | 52 | 0.2 | 8.5 | 21.25 |
| 53 | 53 | 0.5 | 5.64 | 14.09 |
| 54 | 54 | 0.5 | 5.60 | 14.01 |
| 55 | 55 | 0.01 | 3.40 | 8.50 |
| 56 | 56 | 0.2 | 8.5 | 21.25 |
| 57 | 57 | 0.2 | 8.5 | 21.25 |
| 58 | 58 | 0.2 | 8.5 | 21.25 |
| 59 | 59 | 0.5 | 4.60 | 11.51 |
| 60 | 60 | 0.5 | 4.07 | 10.18 |

**Supplementary Table S2:** Bond parameters for the native rpL5 peptide.

| Bond type | $K$ (kcal/mol/Å$^2$) | $r_0$ (Å) |
|---|---|---|
| 1 | 60000 | 24.17 |
| 2 | 60000 | 27.43 |
| 3 | 60000 | 20.63 |
| 4 | 60000 | 24.50 |
| 5 | 7.11 | 3.70 |
| 6 | 0.64 | 4.63 |
| 7 | 0.35 | 4.69 |
| 8 | 0.72 | 6.68 |
| 9 | 2.18 | 5.28 |
| 10 | 2.03 | 4.40 |
| 11 | 1.08 | 4.69 |
| 12 | 0.96 | 5.24 |
| 13 | 1.47 | 5.31 |
| 14 | 1.07 | 5.24 |
| 15 | 1.39 | 5.44 |
| 16 | 0.87 | 4.49 |
| 17 | 10.47 | 3.65 |
| 18 | 2.16 | 4.78 |
| 19 | 0.71 | 5.85 |
| 20 | 0.97 | 5.12 |
| 21 | 1.04 | 5.03 |
| 22 | 1.28 | 5.07 |

| | | |
|---|---|---|
| 23 | 2.31 | 4.65 |
| 24 | 1.70 | 4.87 |
| 25 | 1.28 | 4.81 |
| 26 | 2.40 | 5.38 |
| 27 | 1.67 | 5.44 |
| 28 | 1.51 | 5.26 |
| 29 | 0.72 | 5.77 |
| 30 | 0.44 | 5.03 |
| 31 | 6.16 | 4.03 |
| 32 | 7.61 | 3.84 |
| 33 | 1.96 | 5.02 |
| 34 | 1.61 | 5.61 |
| 35 | 1.35 | 5.25 |
| 36 | 2.30 | 4.31 |
| 37 | 2.26 | 4.73 |
| 38 | 1.92 | 4.98 |
| 39 | 1.78 | 4.84 |
| 40 | 1.41 | 5.25 |
| 41 | 1.39 | 5.19 |
| 42 | 1.50 | 5.17 |
| 43 | 1.79 | 5.29 |
| 44 | 0.57 | 7.32 |
| 45 | 0.56 | 6.88 |
| 46 | 0.46 | 6.97 |
| 47 | 0.68 | 5.91 |
| 48 | 1.31 | 4.67 |
| 49 | 1.74 | 4.57 |
| 50 | 1.42 | 5.10 |
| 51 | 2.04 | 4.53 |
| 52 | 0.56 | 5.86 |
| 53 | 0.20 | 5.61 |
| 54 | 0.34 | 4.89 |
| 55 | 1.02 | 5.48 |
| 56 | 0.72 | 6.56 |
| 57 | 0.70 | 6.51 |
| 58 | 0.75 | 6.06 |
| 59 | 2.38 | 4.47 |

**Supplementary Table S3:** Angle parameters for the native rpL5 peptide.

| Angle type | $K$ (kcal/mol) | $\theta_0$ (degs) |
|---|---|---|
| 1 | 6000000 | 38.08 |
| 2 | 6000000 | 25.58 |
| 3 | 6000000 | 11.84 |
| 4 | 6000000 | 44.53 |

| | | |
|---|---|---|
| 5 | 2.90 | 120.66 |
| 6 | 1.46 | 88.20 |
| 7 | 3.21 | 92.11 |
| 8 | 4.47 | 77.38 |
| 9 | 2.47 | 101.29 |
| 10 | 2.31 | 113.32 |
| 11 | 5.70 | 93.42 |
| 12 | 3.42 | 67.41 |
| 13 | 7.12 | 98.43 |
| 14 | 3.62 | 68.99 |
| 15 | 3.95 | 95.38 |
| 16 | 2.45 | 113.15 |
| 17 | 5.48 | 85.38 |
| 18 | 2.50 | 99.63 |
| 19 | 5.60 | 74.87 |
| 20 | 3.33 | 93.10 |
| 21 | 2.71 | 114.72 |
| 22 | 5.06 | 72.89 |
| 23 | 5.74 | 111.86 |
| 24 | 3.55 | 82.63 |
| 25 | 7.93 | 78.78 |
| 26 | 5.32 | 103.35 |
| 27 | 5.42 | 64.24 |
| 28 | 3.66 | 96.05 |
| 29 | 2.09 | 106.70 |
| 30 | 4.28 | 111.79 |
| 31 | 4.07 | 80.47 |
| 32 | 3.44 | 94.31 |
| 33 | 3.02 | 95.35 |
| 34 | 2.98 | 121.51 |
| 35 | 5.02 | 73.45 |
| 36 | 2.89 | 114.27 |
| 37 | 4.23 | 78.12 |
| 38 | 2.67 | 101.09 |
| 39 | 4.43 | 87.34 |
| 40 | 3.07 | 98.51 |
| 41 | 4.48 | 78.73 |
| 42 | 3.03 | 81.96 |
| 43 | 4.99 | 72.83 |
| 44 | 3.57 | 89.60 |
| 45 | 1.70 | 130.68 |
| 46 | 4.49 | 80.25 |
| 47 | 2.50 | 109.36 |
| 48 | 2.14 | 85.62 |
| 49 | 2.89 | 84.48 |

| | | |
|---|---|---|
| 50 | 2.78 | 77.81 |
| 51 | 1.66 | 112.64 |
| 52 | 5.91 | 67.05 |
| 53 | 2.85 | 96.20 |
| 54 | 4.34 | 73.48 |
| 55 | 2.54 | 97.07 |

**Supplementary Table S4:** Quadratic dihedral parameters for the native rpL5 peptide.

| Dihedral type | $K$ (kcal/mol) | $\phi_0$ (rads) |
|---|---|---|
| 1 | 6000000 | 0.7604 |
| 2 | 6000000 | 0.3260 |
| 3 | 6000000 | 2.1276 |
| 51 | 6000000 | 2.5908 |
| 52 | 6000000 | 2.2405 |
| 53 | 6000000 | 2.1536 |

**Supplementary Table S5:** Class2 dihedral parameters for the native rpL5 peptide. The mbt, ebt, at, aat, and bb13 parameters are all set to 0.

| Dihedral type | $K_1$ (kcal/mol) | $\phi_{01}$ (degs) | $K_2$ (kcal/mol) | $\phi_{02}$ (degs) | $K_3$ (kcal/mol) | $\phi_{03}$ (degs) |
|---|---|---|---|---|---|---|
| 4 | 0.35 | 192.09 | 0.17 | -36.38 | 0.05 | 209.29 |
| 5 | 0.25 | 88.11 | 0.38 | 94.00 | 0.00 | 92.25 |
| 6 | 0.90 | 202.93 | 0.04 | -39.07 | 0.03 | 150.19 |
| 7 | 1.17 | -3.08 | 0.21 | 171.27 | 0.00 | 102.96 |
| 8 | 0.19 | 51.01 | 0.37 | 160.82 | 0.05 | 253.04 |
| 9 | 0.84 | 174.34 | 0.54 | 45.49 | 0.00 | 83.18 |
| 10 | 0.60 | 201.82 | 0.22 | 108.45 | 0.09 | 114.99 |
| 11 | 1.01 | 144.54 | 0.16 | 97.05 | 0.00 | 93.28 |
| 12 | 1.21 | 204.55 | 0.17 | 140.09 | 0.00 | 82.42 |
| 13 | 0.37 | 129.81 | 0.13 | 107.51 | 0.00 | 95.26 |
| 14 | 0.73 | 216.64 | 0.32 | 99.59 | 0.00 | 91.89 |
| 15 | 0.90 | 252.26 | 0.23 | 134.04 | 0.33 | 191.76 |
| 16 | 0.41 | 150.27 | 0.29 | 94.02 | 0.00 | 111.68 |
| 17 | 0.21 | 199.40 | 0.49 | 80.89 | 0.05 | 0.95 |
| 18 | 0.78 | 236.42 | 0.60 | 143.37 | 0.25 | 155.83 |
| 19 | 0.68 | 197.31 | 0.25 | 76.18 | 0.08 | 171.15 |
| 20 | 0.99 | 111.89 | 0.06 | 44.15 | 0.00 | 106.72 |
| 21 | 1.18 | 185.92 | 0.15 | 42.63 | 0.05 | 73.28 |
| 22 | 0.87 | 5.23 | 0.64 | 141.11 | 0.12 | 197.37 |
| 23 | 1.40 | 152.90 | 0.32 | 40.89 | 0.00 | 89.92 |
| 24 | 1.31 | 192.95 | 0.13 | 102.51 | 0.05 | 124.68 |
| 25 | 0.56 | 76.81 | 0.34 | 86.04 | 0.18 | 171.77 |

| | | | | | | |
|---|---|---|---|---|---|---|
| 26 | 0.74 | 133.14 | 0.17 | 80.98 | 0.00 | 92.05 |
| 27 | 0.68 | 183.81 | 0.22 | 15.86 | 0.03 | 63.88 |
| 28 | 0.28 | 15.57 | 0.40 | 152.34 | 0.01 | 186.48 |
| 29 | 0.14 | -19.80 | 0.44 | 147.16 | 0.02 | 22.56 |
| 30 | 0.77 | 181.30 | 0.22 | 120.81 | 0.00 | 185.70 |
| 31 | 0.99 | 193.81 | 0.26 | 85.19 | 0.01 | 114.23 |
| 32 | 0.74 | 154.43 | 0.16 | 75.64 | 0.00 | 85.07 |
| 33 | 0.97 | 188.73 | 0.15 | 91.70 | 0.00 | 141.62 |
| 34 | 0.18 | 143.51 | 0.21 | 160.09 | 0.05 | 12.22 |
| 35 | 0.26 | 200.66 | 0.36 | 137.48 | 0.09 | 237.57 |
| 36 | 0.00 | 99.42 | 0.39 | 118.04 | 0.14 | 83.38 |
| 37 | 0.00 | 92.68 | 0.59 | 163.61 | 0.43 | 117.77 |
| 38 | 0.39 | -55.81 | 0.16 | 84.16 | 0.08 | 102.00 |
| 39 | 0.81 | 193.10 | 0.20 | 162.19 | 0.00 | 228.93 |
| 40 | 0.41 | 184.88 | 0.04 | 140.44 | 0.01 | 0.97 |
| 41 | 0.55 | 158.94 | 0.06 | 94.56 | 0.06 | 231.32 |
| 42 | 0.40 | 184.10 | 0.35 | 66.24 | 0.00 | 29.64 |
| 43 | 0.59 | 204.96 | 0.33 | 104.71 | 0.07 | 140.42 |
| 44 | 0.19 | 162.15 | 0.29 | 102.99 | 0.04 | 139.83 |
| 45 | 0.28 | 198.62 | 0.37 | 90.11 | 0.20 | 119.32 |
| 46 | 0.24 | 190.76 | 0.19 | 90.03 | 0.11 | 79.27 |
| 47 | 0.00 | 98.76 | 0.30 | 83.13 | 0.15 | 110.90 |
| 48 | 0.40 | 23.30 | 0.32 | 134.82 | 0.05 | 159.42 |
| 49 | 0.28 | -36.25 | 0.36 | 115.95 | 0.03 | -2.61 |
| 50 | 0.29 | 175.94 | 0.23 | 110.73 | 0.00 | 64.62 |

**Supplementary Table S6:** Modified LJ parameters for the A0 N130 mutant. The rest of the parameters are the same as the native peptide given in Supplementary Tables 1-5.

| Type 1 | Type 2 | $\epsilon$ (kcal/mol) | $\sigma$ (Å) | $r_c$ (Å) |
|---|---|---|---|---|
| 6 | 6 | 0.25 | 3.39 | 8.47 |
| 7 | 7 | 0.25 | 3.39 | 8.47 |
| 10 | 10 | 0.25 | 3.39 | 8.47 |
| 12 | 12 | 0.25 | 3.39 | 8.47 |
| 17 | 17 | 0.25 | 3.39 | 8.47 |

**Supplementary Table S7:** Modified LJ parameters for the A1 N130 mutant. The rest of the parameters are the same as the native peptide given in Supplementary Tables 1-5.

| Type 1 | Type 2 | $\epsilon$ (kcal/mol) | $\sigma$ (Å) | $r_c$ (Å) |
|---|---|---|---|---|
| 20 | 20 | 0.25 | 3.39 | 8.47 |
| 22 | 22 | 0.25 | 3.39 | 8.47 |
| 24 | 24 | 0.25 | 3.39 | 8.47 |
| 25 | 25 | 0.25 | 3.39 | 8.47 |

| | | | | |
|---|---|---|---|---|
| 27 | 27 | 0.25 | 3.39 | 8.47 |

**Supplementary Table S8:** Modified LJ parameters modified for the A2 N130 variant. The rest of the parameters are the same as the native peptide given in Supplementary Tables 1-5.

| Type 1 | Type 2 | $\epsilon$ (kcal/mol) | $\sigma$ (Å) | $r_c$ (Å) |
|---|---|---|---|---|
| 33 | 33 | 0.25 | 3.39 | 8.47 |
| 34 | 34 | 0.25 | 3.39 | 8.47 |
| 35 | 35 | 0.25 | 3.39 | 8.47 |
| 37 | 37 | 0.25 | 3.39 | 8.47 |
| 39 | 39 | 0.25 | 3.39 | 8.47 |
| 40 | 40 | 0.25 | 3.39 | 8.47 |
| 41 | 41 | 0.25 | 3.39 | 8.47 |
| 42 | 42 | 0.25 | 3.39 | 8.47 |
| 43 | 43 | 0.25 | 3.39 | 8.47 |
